# PEGylated surfaces for the study of DNA-protein interactions by atomic force microscopy

**DOI:** 10.1101/680561

**Authors:** Bernice Akpinar, Nicholas A. W. Bell, Alice L.B. Pyne, Bart W. Hoogenboom

## Abstract

DNA-protein interactions are vital to cellular function, with key roles in the regulation of gene expression and genome maintenance. Atomic force microscopy (AFM) offers the ability to visualize DNA-protein interactions at nanometre resolution in near-physiological buffers, but it requires that the DNA be adhered to the surface of a solid substrate. This presents a problem when working at biologically relevant protein concentrations, where protein may be present at large excess in solution; much of the biophysically relevant information can therefore be occluded by non-specific protein binding to the underlying substrate. Here we explore the use of PLL_x_-*b*-PEG_y_ block copolymers to achieve selective adsorption of DNA on a mica surface. Through varying both the number of lysine and ethylene glycol residues in the block copolymers, we show selective adsorption of DNA on mica that is functionalized with a PLL_10_-*b*-PEG_113_ / PLL_1000-2000_ mixture as viewed by AFM imaging in a solution containing high concentrations of streptavidin. We show that this selective adsorption extends to DNA-protein complexes, through the use of biotinylated DNA and streptavidin, and demonstrate that DNA-bound streptavidin can be unambiguously distinguished by in-liquid AFM in spite of an excess of unbound streptavidin in solution.

## Introduction

Interactions between DNA and proteins regulate a number of processes crucial to cellular function that include transcription, chromosome maintenance, DNA replication and repair. DNA-binding proteins employ a range of different mechanisms to interact with both select and non-select sites on DNA.^1^ Key mechanistic insights have been revealed using biophysical techniques such as fluorescence microscopy,^2–4^ optical tweezers,^5,6^ surface plasmon resonance,^7^ and atomic force microscopy (AFM).^8,9^

AFM has been established as a powerful single-molecule technique to probe DNA-protein interactions, due to its ability to directly image DNA at nanometre resolution in physiologically relevant conditions without the need for labelling.^10^ However, to obtain high-resolution images of biomolecules in liquid, the sample must be adhered to an underlying solid support. Muscovite mica is the substrate of choice for AFM imaging of DNA, due to the ease of preparing an atomically flat mica surface via cleavage along the basal plane, and due to the polar, hydrophilic nature of the cleaved surface, facilitating the adsorbing and retention of biomolecules in aqueous solution. When mica is hydrated, K^+^ ions dissociate from interstitial sites within mica’s aluminium phyllosilicate lattice, resulting in a net negative charge on the surface. To permit the adsorption of DNA with its highly negatively charged phosphate backbone, the negative surface charge needs to be screened or compensated for.^11,12^ Many adsorption protocols have been established for the 2D confinement of DNA to pre-treated mica,^13^ one of the most commonly adopted being the use of transition metal cations such as Ni^2+^, Co^2+^and Zn^2+^ that can substitute into vacant sites within the mica lattice, yielding positively charged patches for the adsorption of DNA.^14^ The strength of the electrostatic attraction can be modified by the presence of additional ions and chelating agents within imaging buffer.^15,16^ Other methods to facilitate DNA absorption include modification of the surface chemistry using silanes,^13^ the formation of partially positively charged lipid bilayers^9,17^ and the electrostatic adsorption of positively charged polymers such as poly-L-lysine (PLL).^18,19^

The aforementioned approaches are often adopted for the study of DNA-protein binding using AFM. However, they can result in non-specific protein-surface interactions, which are non-trivial to deconvolute from specific DNA-protein interactions. The problem of non-specific adsorption can be addressed by the use of surface coatings that are protein repellent, for example polymer brushes that suppress protein binding by steric repulsion.^20^ One approach to suppress protein binding is to create an interfacial layer of polyethylene glycol (PEG) brushes. The high degree of hydration and flexibility of these brushes causes surface passivation when the chains are of sufficient length and grafted at high density.^21^ Facile preparation of PEGylated surfaces is achieved using multifunctional copolymers comprising both surface binding domains and surface passivating PEG domains. Graft-copolymers with a cationic PLL backbone and PEG side chains (PLL-*g*-PEG) have proven particularly effective at self-assembling into densely packed polymeric brushes to form non-fouling surfaces.^22–24^ In addition, bio-recognition sites, such as RGD-peptides, have been incorporated into these films to promote cell adhesion whilst suppressing the non-specific adsorption of serum proteins.^25,26^ Similarly, the incorporation of biotin-terminated PEG chains has been used to form small molecule biosensors that selectively bind streptavidin, neutravidin and avidin.^27^ Unmodified PLL-*g*-PEG films have also shown the ability to selectively adsorb DNA polyelectrolytes to the underlying positively charged PLL layer, whilst the PEG layer remains impervious to other proteins, as confirmed by fluorescence imaging.^28^

The well-studied graft copolymer (PLL-*g*-PEG) adopts a comb-like conformation in solution comprised of a long PLL backbone with randomly distributed PEG side chains, whilst the block copolymer (PLL-*b*-PEG) exhibits a linear worm-like conformation comprised of regions of lysine repeats followed by regions of ethylene glycol repeats. Both copolymers can form protein repellent PEG brushes on a variety of substrates through the spontaneous electrostatic attachment of their lysine residues. In the case of the graft copolymer, the length of the PEG block and the grafting ratio affect the density and hence efficacy of the anti-fouling brushes.^24^ The diblock copolymer has been less widely employed for surface passivation but has been shown to be effective at inhibiting cell adhesion on glass surfaces micro patterned with PLL_100_-*b*-PEG_22_.^29^ The passivation properties of the diblock copolymer (PLL_x_-*b*-PEG_y_) can be tuned by varying the degree of polymerization of both the PLL (x) and PEG (y) chains which would affect the packing density onto the underlying substrate, although as of yet variations of these have not been explored. We here set out to determine whether linear PLL_x_-*b*-PEG_y_ diblock copolymers can be used in the functionalization of mica to yield a surface that selectively adsorbs DNA and allows characterization of DNA-protein complexes by AFM. Through optimization of the composition of the diblock copolymer, we have developed biphasic films which promote the adsorption of negatively charged DNA, whilst passivating against non-specific protein adsorption. Specifically, we perform mica surface functionalization which allows high-resolution AFM imaging of DNA and of DNA-protein complexes in solution whist resisting non-specific protein adsorption.

## Methods

### Materials

Relaxed plasmid pBR322 DNA was purchased from Inspiralis Ltd. A 672bp length of DNA was prepared by PCR amplification of a section of lambda DNA (New England Biolabs) using a forward primer 5’-CGATGTGGTCTCACAGTTTGAGTTCTGGTTCTCG-3 and reverse primer 5’-GGAAGAGGTCTCTTAGCGGTCAGCTTTCCGTC-3’ purchased from Integrated DNA Technologies. Each primer was labelled at its 5’ end with a single biotin thereby resulting in a double-stranded DNA PCR product labelled at both ends with biotin. The PCR product was purified using a QIAquick PCR purification kit (Qiagen). Monovalent streptavidin was produced by the Howarth lab.^30^ Block copolymers methoxy-poly(ethylene glycol)-block-poly(L-lysine hydrochloride) with varying degrees of polymerization of the poly-L-lysine and polyethylene glycol blocks were purchased as lyophilized powders from Alamanda polymers. The polymers used for this study were PLL_10_–PEG_22_, PLL_10_–PEG_113_, PLL_100_–PEG_113_ and PLL_10_– PEG_454_ where the subscript refers to the degree of polymerization, i.e. the number of monomer repeats. Supplementary table 1 details the corresponding molecular weights for each of the polymers used. A 0.01% w/v solution of poly-L-lysine (PLL_1000-2000_) with approximately one HBr molecule per lysine residue, along with all other reagents, were purchased from Sigma-Aldrich.

### Agarose gel electrophoresis

Biotinylated DNA binding to mono and tetravalent DNA was verified by AGE (1% agarose, 1 × TBE) using the BioRad Wide Mini-Sub Cell GT Electrophoresis System. 5□μl of pre-incubated samples were mixed with 1□μl of 6 × loading buffer before loading onto the agarose gel. The samples were allowed to migrate for 90 minutes (running buffer: 1 × TBE; 90□V). The gel was stained for 40 minutes in a solution 3 × GelRed (Biotium) and visualized using UV light.

### Mica modification and DNA deposition

For the preparation of copolymer films, freshly cleaved mica discs (diameter: 5 mm) were covered in 10 μl of solution comprising only PLL_x_-b-PEG_y_ (1 mg/ml in MilliQ water) or a mixture of 5 μl of the PLL_x_-b-PEG_y_ solution and 5 μl PLL_1000-2000_ (0.01 % w/v). Mica discs were incubated with these solutions for 45 minutes in a humid environment, before washing 5 times with MilliQ water and 5 times with imaging buffer (10 mM phosphate buffer pH 7.4). 5 μl of DNA plasmid (∼1.5 ng/μl) or 3 μl biotinylated DNA (∼3.5 nM) was immediately added to the disc and allowed to equilibrate for approximately 10 minutes prior to imaging. A similar protocol was followed for functionalization with PLL alone but the PLL incubation time was reduced to 1 minute before thoroughly rinsing to minimize surface contamination. A solution of PLL, either 0.01 % (Fig.3(a)) or 0.001 % (Fig.2) was used to form full or partial monolayers, onto which DNA could be adsorbed.

**Figure 1.**
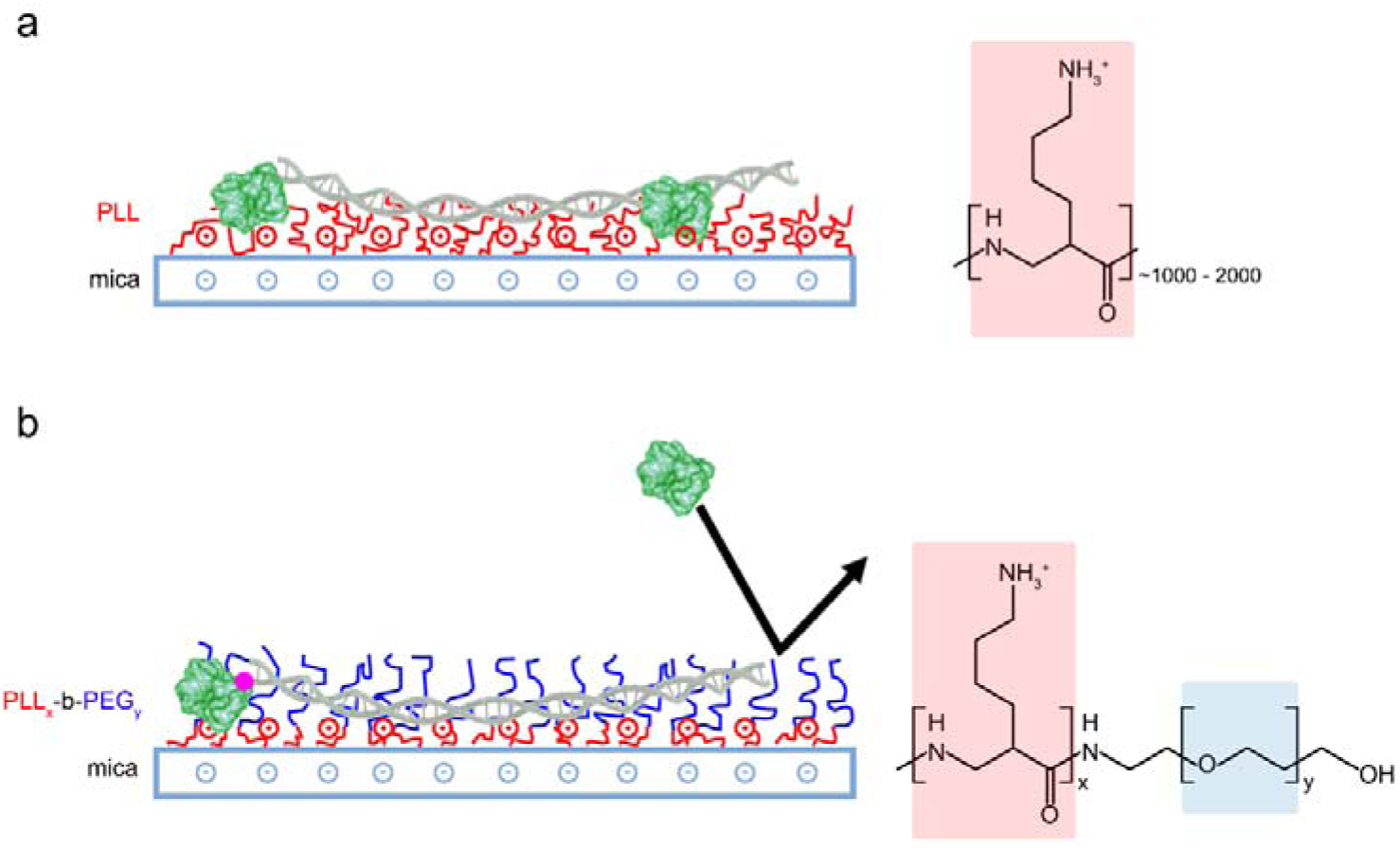
Schematic representation of different DNA surface adsorption methods. (a) Adsorption of DNA and proteins is promoted by modifying mica substrates with positively charged long-chain PLL_1000-2000_. (b) PLL_x_-b-PEG_y_ block copolymers form films of densely packed PEG chains that repel proteins whilst the accessible lysine residues promote electrostatic adsorption of the highly charged DNA only.

**Figure 2.**
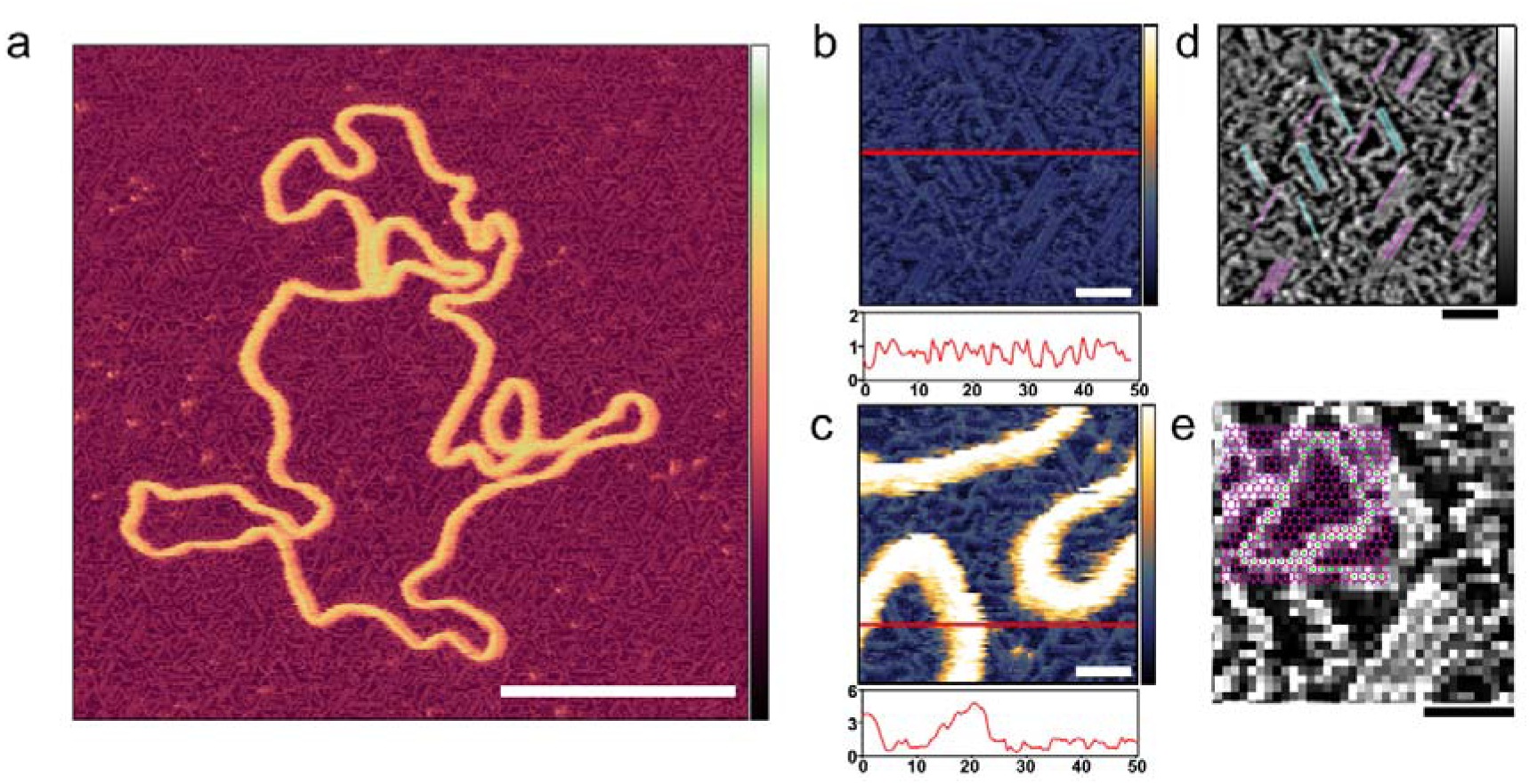
Ordering of poly-L-lysine chains on a mica substrate. (a) AFM image with tip sampling every ∼0.5 nm showing a DNA plasmid adsorbed onto PLL_1000-2000_-functionalized mica. (b,c) At higher magnification, PLL chains are unambiguously resolved. Height profiles underneath provide an indication of the respective protrusions of the PLL chains and of the DNA. (d) the axis of alignment observed in (b) are highlighted (e) the mica lattice geometry is here aligned and overlaid with the resolved lysine chains, their corresponding overlap with vacancies is observed. Inset colour scale for (a) is 8 nm and inset colour scales for (b) and (c) 4 nm, for (d) and (e) 0.8 nm. Scale bar for (a) is 100 nm, 10 nm for (b), (c) and (d).

**Figure 3.**
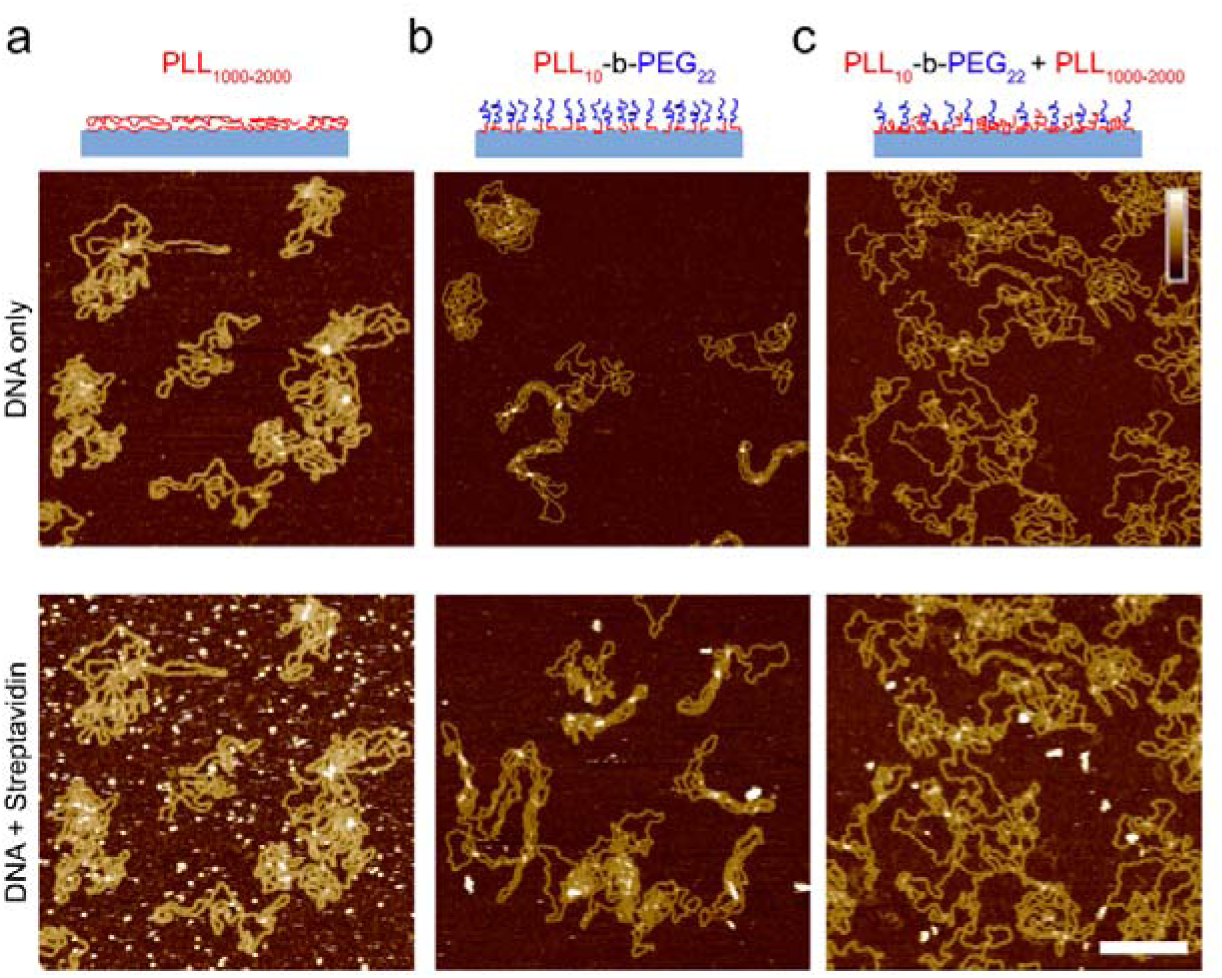
Characterization of the adsorption of DNA plasmid and streptavidin on functionalized mica. Streptavidin (160 nM) was added after DNA immobilization, DNA was incubated for 10 minutes prior to imaging and streptavidin was left to equilibrate for 10 minutes prior to imaging on (a) PLL_1000-2000_ only surface, (b) PLL_10_-b-PEG_22_ and (c) a mixed PLL_10_-b-PEG_22_/PLL_1000-2000_ surface. Colour scale (inset in c) 10 nm; scale bar 200 nm.

### AFM imaging

All AFM imaging was carried out at room temperature with the samples hydrated in imaging buffer. Data were recorded using a Dimension FastScan Bio AFM (Bruker, Santa Barbara, USA) operated in PeakForce Tapping mode. Force-distance curves were recorded over 10-40 nm (PeakForce Tapping amplitude of 5-20 nm), at a frequency of 8 kHz. FastScan D (Bruker) cantilevers were used for all imaging (resonance frequency of ∼110 kHz, nominal spring constant ∼0.25 Nm^−1^). Images were processed using first-order line-by-line flattening and zeroth order plane fitting to remove sample tilt and background using Gwyddion.

### Quantification of protein binding by AFM

To quantify the amount of background protein in the experiments using DNA plasmids, AFM images were thresholded using Gwyddion, to differentiate streptavidin molecules (∼4 nm height), DNA molecules (∼2 nm height), and the substrate (∼0 nm height), with the height thresholds adjusted to give the best identification of streptavin and DNA, as determined by visual inspection. For each quantification, a total area of at least 74 μm^2^ was analyzed to give the percentage surface coverage of streptavidin shown in Fig.4(c). In the studies of the biotinylated DNA, individual streptavidin molecules were counted using ImageJ to obtain the number of streptavidin molecules that were not bound to the ends of DNA, with results shown in Fig. S3.

**Figure 4.**
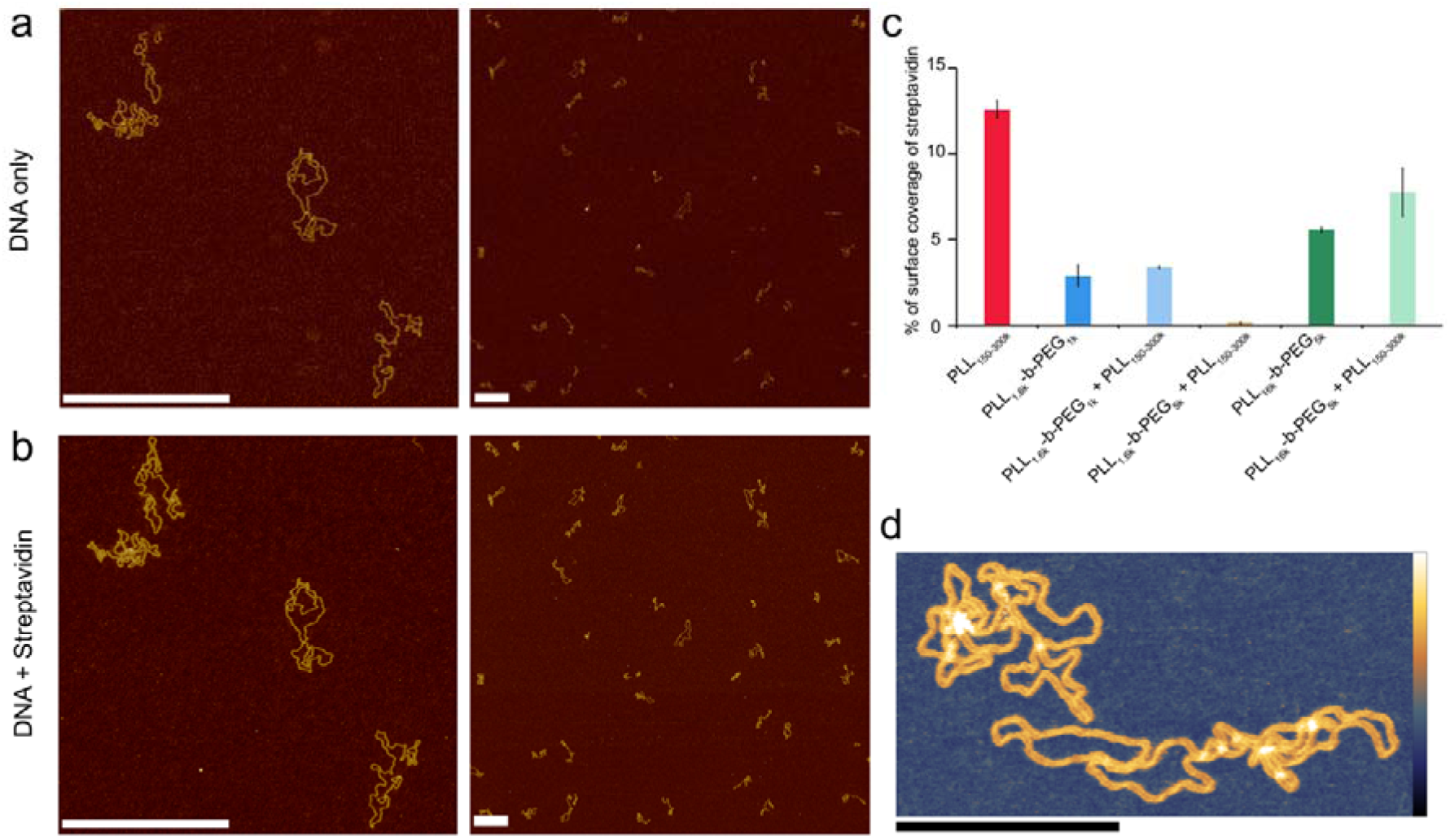
Optimized Poly(L-lysine)-b-Poly(ethylene glycol) surfaces for exclusive DNA adsorption. AFM images showing selective DNA adsorption on PLL_10_-b-PEG_113_ / PLL_1000-2000_ surfaces. (a) DNA plasmid only. (b) The same area following the addition of 160 nM streptavidin. (c) Percentage background streptavidin coverage at 160 nM for functionalization protocols using different PEG chain lengths, error bars correspond to the minimum and maximum values as determined from two different areas. (d) A higher resolution image of DNA on the PLL_10_-b-PEG_113_ / PLL_1000-2000_ surface. Colour scales (see inset in Fig. 2(a)) for (a) and (b) are 7 nm, (d) with inset colour scale is 9 nm. Scale bars in (a) and (b) are 500 nm and in (d) 200 nm.

## Results and Discussion

Poly-L-lysine (PLL) has been used extensively to immobilize DNA on a mica surface. This immobilization depends on an interfacial layer of positively charged long-chain (PLL) polymers that bind DNA through electrostatic attraction. However, this surface also facilitates the non-specific adsorption of proteins, thus complicating the identification of targeted DNA-protein interactions. Copolymers comprising both PLL and PEG repeats achieve reduced non-specific protein adsorption through an additional PEG component, which acts as a steric barrier to protein binding (Fig. 1). To compare, on one hand, the efficiency of diblock PLL_x_-b-PEG_y_ copolymers for specific DNA and DNA-protein immobilization, and, on the other hand, immobilization using traditional PLL protocols, we first characterized DNA adsorption on a mica functionalized with PLL only. PLL spontaneously attaches to the negatively charged mica surface via its highly charged lysine residues (pKa ∼ 10.5) to yield homopolymer films stable over a range of pH and salt concentrations.^31^ The PLL used here was PLL_1000-2000_ where the subscript refers to the number of lysine repeats in the homopolymer, corresponding to a molecular weight of 150,000-300,000 g mol^−1^. See ESI (table S1) for molecular weights of the other polymers used in this study.

PLL surface functionalization is obtained by incubation of a cleaved mica surface in PLL solution. Deposition at low concentrations (0.001% w/v), allows relaxation of the lysine chains and adsorption in flattened, stretched out conformations where individual poly-l-lysine molecules are resolved (Fig. 2).^31^ High resolution on the individual lysine chains was achieved in 10 mM phosphate buffer, a relatively poor solvent for poly-L-lysine, reducing repulsion of the AFM tip which can then contact the collapsed chains (Fig. 2(b)).^32^ The PLL chains are seen to preferentially align along three orientations, with an angular difference of about 60° (Fig. 2(d)). PLL chains appear better resolved when aligned at larger angles with respect to the fast scan axis (left to right in these images). When aligned along fast scan axis itself, PLL chains are more difficult to resolve as their width is approximately equal to the width of one scan line (0.5 nm) and therefore more sensitive to the precise position of consecutive lines. The observed orientations are consistent with a molecular arrangement in which the lysine residues occupy interstitial sites on the mica lattice vacated by K^+^ ions of similar van der Waals radii (320 and 280 pm respectively) (Fig. 2(e)).

When deposited from the stock solution at high PLL concentration (0.01% w/v), the lysine chains adopt more globular forms, resulting in an apparently more homogeneous surface functionalization (Fig. 3(a)). PLL functionalized mica enhances the adsorption of DNA, but also of other biomolecules that may be present in solution, including those of reduced charge. This is demonstrated by the immobilization of both the highly negatively charged plasmid DNA and the slightly negatively charged streptavidin (pKa ∼5.0-6.0, at pH 7.4), (Fig. 3(a)).^33^ We show that the surface can be modified to achieve a more preferential, selective adsorption of DNA by functionalization with PLL_10_-b-PEG_22_ alone or by a combination of PLL_10_-b-PEG_22_ and long-chain PLL_1000-2000_. In the presence of the block copolymer streptavidin adheres as sparse clusters (Fig. 3(b)(c)), perhaps due to heterogeneous surface coverage of the protein-repellent PEG. In both cases, large areas of functionalized mica are visible without protein adsorbed. This can be explained by the effective repulsion that arises when the polyethylene-glycol chains form a sufficiently dense steric barrier.

To achieve a homogeneous surface that resists protein adsorption across the entire sample, the PEG molecules should be grafted at a density that is large enough to facilitate overlap between different chains, resulting in the formation of a dense polymer brush.^34^ This requires the radius of gyration *R*_*G*_ for the polymer to be comparable or larger than the mean distance between grafting sites. It follows that longer PEG chains are more effective in passivating a surface against protein binding, provided that they are grafted at sufficient densities.^35^ By increasing the PEG block length (*y)* in the PLL_x_-b-PEG_y_ diblock copolymer, we generated an optimized surface functionalization which resisted non-specific protein adsorption in 160 nM streptavidin (Fig. 4). PLL_10_-b-PEG_113_ constructs were more effective than PLL_10_-b-PEG_22_ in preventing streptavidin binding, however co-functionalization with PLL_1000-2000_ was required to facilitate the adsorption and imaging of DNA; functionalization with the block copolymer alone yielded a densely packed surface that did not appear to bind DNA (data not shown). Finally, for even longer PEG chains (PLL_10_-b-PEG_454_), DNA adsorption appeared to be prevented altogether, even in the additional presence of PLL_1000-2000_ chains (Fig. S1). We cannot exclude that DNA absorption to the underlying PLL layer is obscured by the PEG layer: the hydrodynamic radius of the PEG_454_ is ∼13.7 nm and therefore the film thickness is expected to be much greater than the height of the DNA.

In addition to varying the PEG block length, we studied the effect of changing the PLL chain length × in the diblock copolymer PLL_x_-b-PEG_113_ (Fig. S1). In contrast to PLL_10_-b-PEG_113_, PLL_100_-b-PEG_113_ facilitated DNA adsorption without additional long chain PLL, however this surface was less selective, binding increased streptavidin. This implies that the longer lysine block (in the PLL_x_-b-PEG_113_) increases the spacing between the passivating PEG moieties. In this case the effective grafting distance between these moieties becomes larger than their extension (radius of gyration), such that they adopted collapsed coil conformations and no longer form an effective steric barrier.^20^ We also note that with the increased length PLL in the block copolymer, the DNA plasmids appear more condensed than on PLL_10_-b-PEG_y_, forming toroid and rod-like structures.^36^

Full quantification of streptavidin binding is complicated when considering images with complex arrangements of DNA and streptavidin on the surface. However, PLL_10_-b-PEG_113_ / PLL_1000-2000_ functionalization emerges as the most effective in suppressing streptavidin binding whilst allowing visualization of DNA by AFM, both by qualitative comparison of the images and by tentative quantification (Fig. 4c). To determine if this functionalization is also effective to study DNA-protein interactions, we created a 672bp length of dsDNA with a single biotin at each end by using PCR amplification with biotinylated primers. Biotin binds to streptavidin with an extremely high affinity with a Kd on the order of femtomolar. These binding partners were chosen for the strong binding affinity of their interaction and relatively low dissociation rate (less than 10% of molecules dissociated after 12 hours at 37°C).^30^ Two streptavidin variants were considered: tetravalent streptavidin and monovalent streptavidin. Although both exhibit a similar binding affinity, monovalent streptavidin has only one functional biotin binding subunit compared to four in wild-type streptavidin. This prevents end-tailing of biotin labelled DNA. The binding of both proteins to the dsDNA construct was confirmed by electrophoretic band shift assay (Fig. S2). DNA incubated with a 50x excess of monovalent streptavidin was adsorbed on the PLL_10_-b-PEG_113_ / PLL_1000-2000_ functionalized mica surface (Fig. 5(a)). DNA molecules with streptavidin bound to both biotinylated ends were specifically adsorbed to the surface (Fig 5(b)). The excess monovalent streptavidin in solution, was not observed at high concentration on the surface, implying good non-specific protein passivation. 40% of adsorbed streptavidin molecules were found at the ends of DNA (*n* = 531). The amount of background streptavidin was much higher in these experiments (i.e., with the biotinylated DNA) than for plasmids immobilized on the same surface (Fig. 4). This increased background may be due to streptavidin binding to biotinylated oligomer contaminants, themselves too small to be detected by the AFM tip. The advantages of PLL_10_-b-PEG_113_ / PLL_1000-2000_ functionalization was further demonstrated by comparison with the traditional PLL_1000-2000_ only functionalization which yielded increased adsorption of non-DNA-bound streptavidin on the surface (Fig. S3).

**Figure 5.**
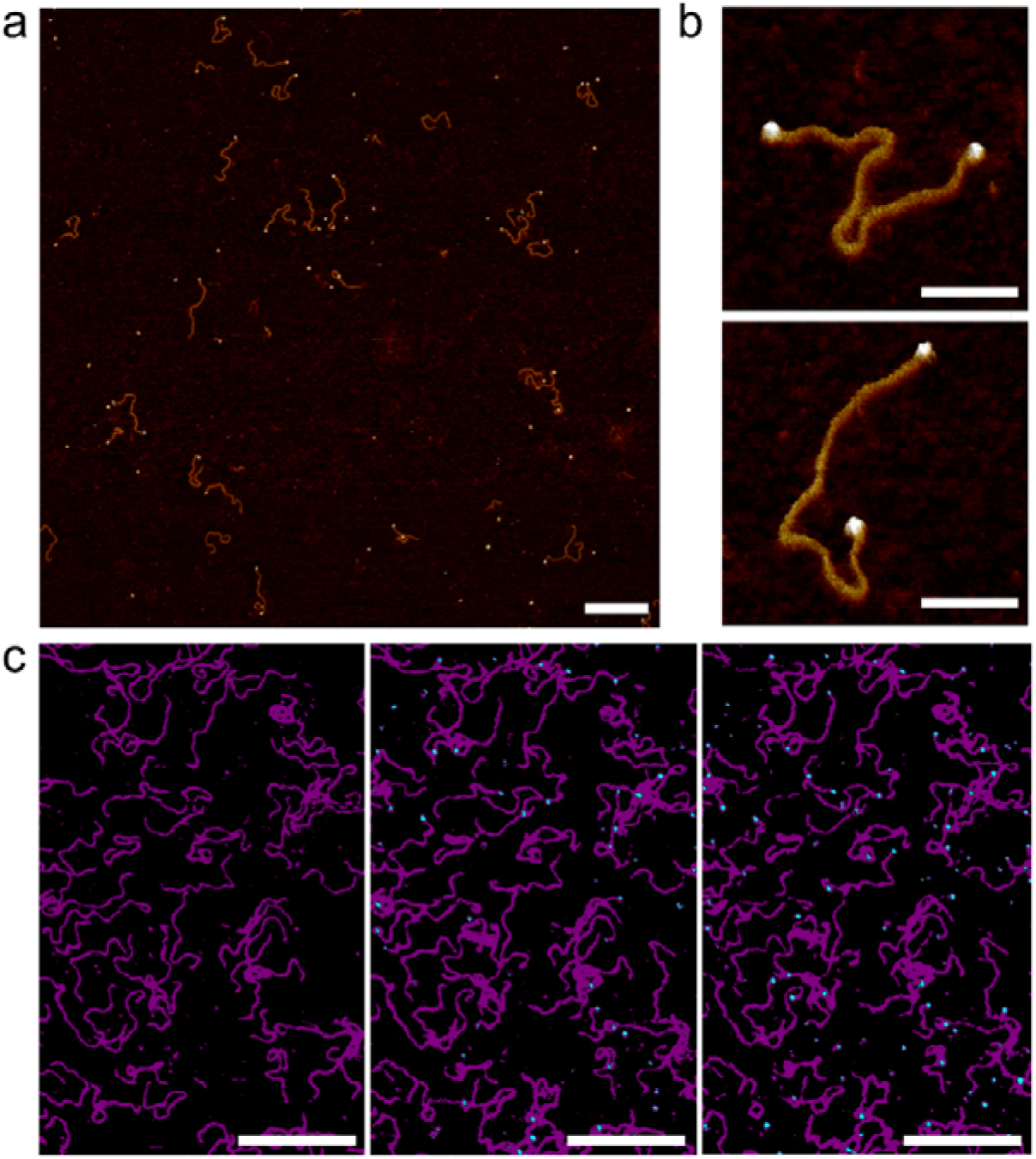
Streptavidin binding to dual-end biotinylated 672 bp DNA on mica treated with PLL_10_-b-PEG_113_ / PLL_1000-2000_. (a) AFM image of 672 bp DNA after pre-incubation with ∼50 molar excess of monovalent mono-streptavidin over biotin tag. (b) High-resolution images showing mono-streptavidin bound to both ends of dual-biotin DNA. (c) An AFM image with the colour scale adjusted to highlight immobilized DNA (magenta) and added tetravalent streptavidin (cyan) for streptavidin concentrations of 0 nM, 150 nM and 750 nM (from left to right). Colour scale (see inset in Fig. 2(a)) is 5 nm. Scale bars for (a) and (c) are 200 nm and for (b) 50 nm.

It is of considerable interest to study the dynamics of DNA-protein interactions. To determine whether this method can used to study the binding of proteins to DNA *in situ*, tetravalent streptavidin was flowed over biotinylated DNA that had already been immobilized on PLL_10_-b-PEG_113_ / PLL_1000-2000_ functionalized mica (Fig. 5(c)). Binding is observed as the formation of cyan protrusions at the ends of the immobilized biotinylated DNA substrates (magenta) which increase from 150 nM to 750 nM. A higher concentration of streptavidin is required for immobilized biotinylated DNA as compared to biotinylated DNA in solution. This suggests limited accessibility of the biotin binding site which is attached to the end of a large DNA molecule and hidden underneath the PEG layer. In this instance tetravalent streptavidin was used as opposed to monovalent streptavidin to increase the surface area for binding and thus reduce steric hinderance effects.^37^ High-resolution AFM imaging requires the surface immobilization of DNA, which can result in the masking of binding sites, and consequently we here found it best to pre-incubate the DNA with the protein prior to surface deposition.

## Conclusions

We have demonstrated the use of hydrophilic diblock copolymers comprising both a cationic surface binding domain (PLL) along with a neutral protein repellent domain (PEG) for the formation of passivating films for the selective immobilization of DNA and DNA-protein complexes. The chain lengths of both blocks were optimized to repel the non-specific adsorption of streptavidin in solution whilst adsorbing highly charged DNA molecules. The surface-passivating properties of this PEG film are demonstrated through the selective binding of biotinylated DNA-streptavidin complexes, minimising non-specific streptavidin surface binding, and could be extended for the study of DNA interactions with other proteins.

## Acknowledgements

The authors thank Richard Thorogate for technical support. This work has been funded the EPSRC studentship (EP/L015277/1 for B.A.); UK EPSRC and MRC fellowships (EP/M5064481 and MR/R024871/1 to A.L.B.P); EPSRC equipment funding (EP/M028100/1).

