## Supplementary Information for "PEGylated surfaces for the study of DNA-protein interactions by atomic force microscopy"

| Notation as degree of polymerization | Notation as molecular weight |
|---|---|
| $PLL_{1000\text{-}2000}$ | $PLL_{150\text{-}300k}$ |
| $PLL_{10}$-b-$PEG_{22}$ | $PLL_{1.6k}$-b-$PEG_{1k}$ |
| $PLL_{10}$-b-$PEG_{113}$ | $PLL_{1.6k}$-b-$PEG_{5k}$ |
| $PLL_{100}$-b-$PEG_{113}$ | $PLL_{16k}$-b-$PEG_{5k}$ |
| $PLL_{10}$-b-$PEG_{454}$ | $PLL_{1.6k}$-b-$PEG_{20k}$ |

**Supplementary Table 1 |** Table expressing molecular weights corresponding to degrees of polymerization for the reagents used in this study.

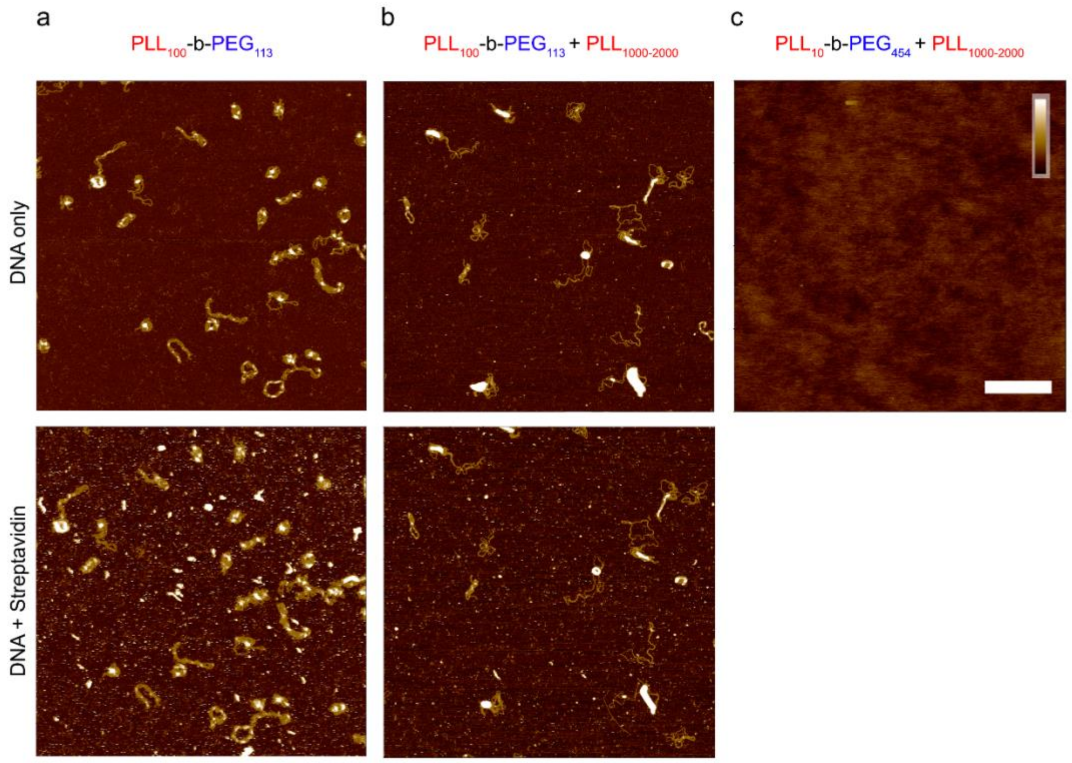


**Supplementary figure 1|Additional characterization of DNA plasmid and streptavidin adsorption on functionalized mica**. Streptavidin (160 nM) was added after DNA immobilization on (a) $PLL_{100}$-b-$PEG_{113}$ surface, (b) a mixed $PLL_{100}$-b-$PEG_{113}$ and $PLL_{1000\text{-}2000}$ surface and (c) $PLL_{100}$-b-$PEG_{454}$ surface. Inset colour scale 8 nm and scale bar 200 nm

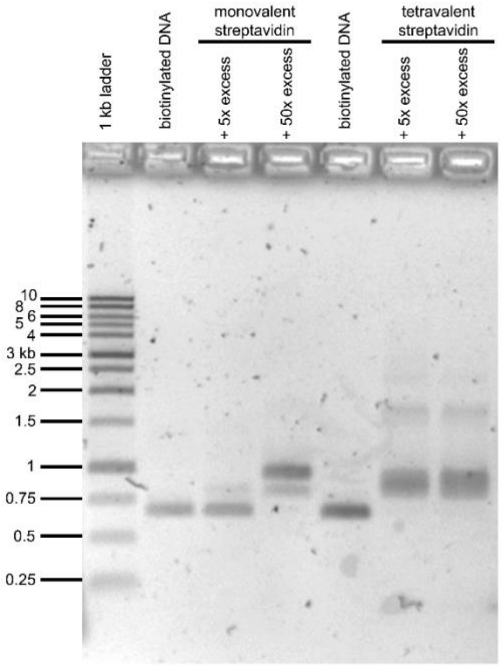


**Supplementary figure 2|Monovalent and tetravalent streptavidin binding to dual-biotin 672 bp DNA.** Clear band shifts confirm binding of monovalent streptavidin when pre-incubated for 5 minutes at ~50x excess and tetravalent streptavidin when at ~5x excess over the number of biotin binding sites.

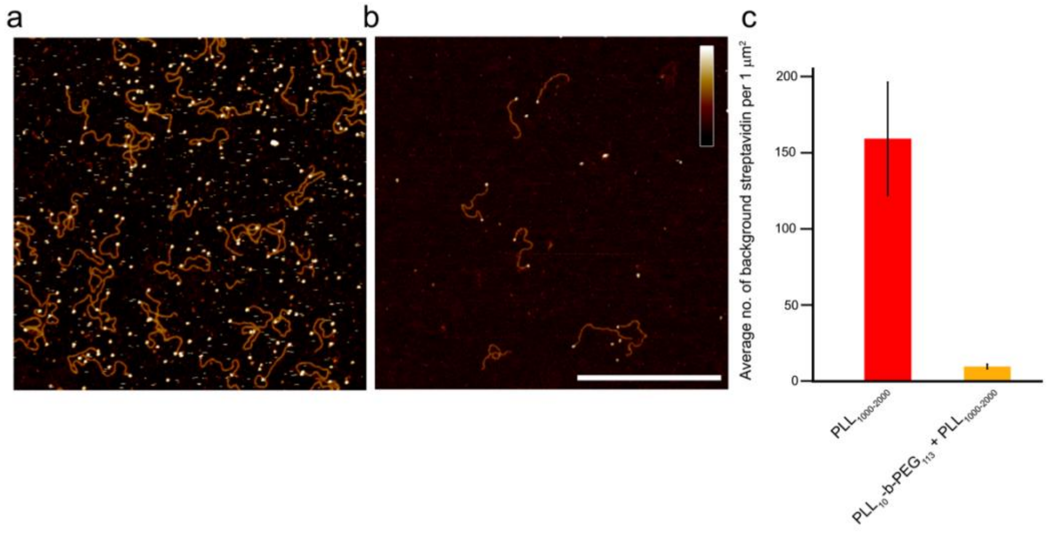


**Supplementary figure 3|Dual-biotin 672 bp DNA pre-incubated with monovalent streptavidin** added to (a) PLL$_{1000\text{-}2000}$ and (b) PLL$_{10}$-b-PEG$_{113}$ / PLL$_{1000\text{-}2000}$ surface. (c) a graph of the average number of streptavidin molecules bound to the background (i.e. not on the end of a DNA molecule) per 1 μm$^2$ analyzed for passivated surface (mean ± standard deviation, $n$ = 8 images, 32 μm$^2$) and for the PLL surface ($n$ = 7 images, 28 μm$^2$). Scale bar is 400 nm and height scale is 6 nm.
